# Super-Resolution Imaging Reveals Stretch-Induced Architectural Rearrangement of Desmoplakin in Desmosomes

**DOI:** 10.1101/2025.08.26.672431

**Authors:** Leslie D. Seeley, Collin M. Ainslie, Mary Kathryn Sewell-Loftin, Alexa L. Mattheyses

## Abstract

Desmosomes (DSMs) are intercellular junctions essential for providing mechanical resilience to tissues, particularly the epidermis. Desmoplakin (DP) is a key DSM protein which anchors plaque proteins to keratins, thereby ensuring tissue integrity under mechanical stress. Clinically, DP mutations impair keratinocyte adhesion and structural integrity, leading to skin fragility disorders. However, how mechanical forces influence DSM architecture is poorly understood. We hypothesized that physiological stretch could alter DP architecture in DSMs. To test this, we subjected normal human epidermal keratinocytes (NHEKs) and DP-knockout human keratinocytes expressing either DPI-mEGFP, DP1a-mEGFP, or DP2-mEGFP to mechanical stretch using the Flexcell system (13% uniaxial strain for 30 minutes). Direct stochastic optical reconstruction microscopy (dSTORM) was used to visualize DP architecture with 20 nm resolution. We found mechanical stretch significantly increased the distance between DP cytoplasmic tails compared to static controls across all cell lines. In contrast, there was no significant change in the N-terminal head domain under stretch, highlighting the tail domain as the primary site of mechanical adaptation. This work enhances our understanding of how DSMs and DP isoforms respond to biomechanical forces, revealing that the C-term of DP undergoes a strain-induced conformational shift, reorganizing the DSM architecture in response to physiological stress. Ultimately, elucidating the spatial and biomechanical behavior of DP will deepen our understanding of its contribution to dermatological health and disease.

## To the editor

Desmosomes (DSMs) are intercellular junctions that confer mechanical resilience to the epidermis. Central to this function is the plakin protein desmoplakin (DP), which anchors the desmosomal plaque to the keratin intermediate filament cytoskeleton. Pathogenic variants in DP impair keratinocyte adhesion and compromise structural integrity, contributing to a spectrum of dermatological disorders, including striate palmoplantar keratoderma, Carvajal syndrome, and lethal acantholytic epidermolysis bullosa (Perl et al., 2024). Affected individuals typically exhibit both cutaneous and extracutaneous manifestations characterized by increased skin fragility, hyperkeratosis, and blistering under mechanical stress, underscoring the importance of investigating DSM biomechanics. Structurally, DSMs have a tripartite organization: transmembrane desmosomal cadherins mediate binding in the extracellular domain (ECD); the outer dense plaque (ODP) contains plakoglobin, plakophilins, and the N-terminal DP head domain; and the inner dense plaque (IDP) includes the DP tail domain, which binds keratin filaments to form a cytoskeletal linkage (Fig. 1a) (North et al., 1999; Russell et al., 2004). This IDP connection supports a tensile filament network that distributes mechanical load and resists forces from stretching, friction, and pressure (Broussard et al., 2020; North et al., 1999).

**Figure 1.**
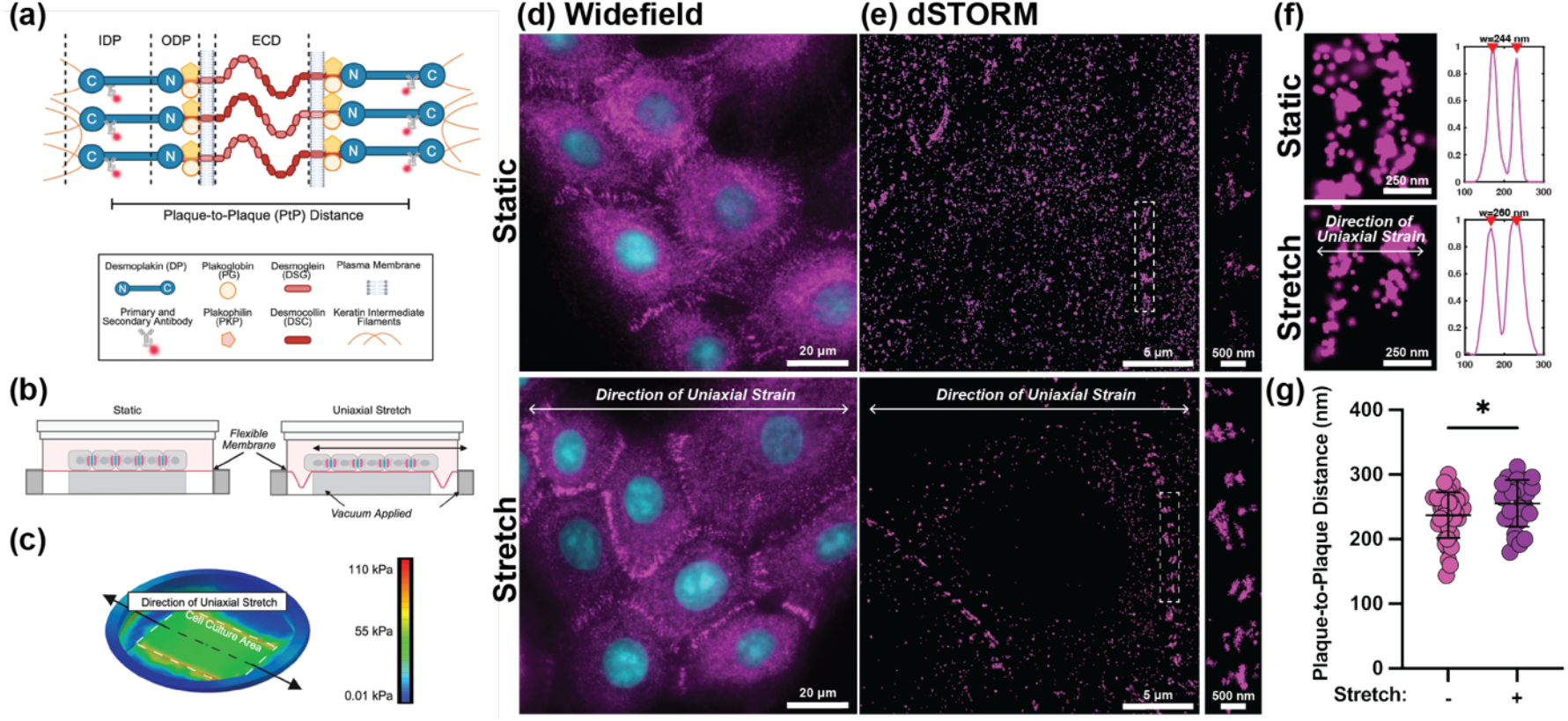
Desmoplakin architecture undergoes nanoscale remodeling in human keratinocytes in response to uniaxial mechanical stretch. (a) Schematic of desmosomal organization highlighting the spatial domains of desmoplakin (DP), associated plaque proteins (plakoglobin (PG), plakophilin (PKP)), and cadherins (desmoglein (DSG), desmocollin (DSC)). (b) Diagram of experimental setup showing NHEKs cultured on flexible membranes under static or stretch conditions using a vacuum-based Flexcell system. (c) Finite element modeling of uniaxial stretch across the cell culture area indicates a uniform applied stress of 55 kPa. (d) Widefield images of DPI (magenta) and DAPI (cyan) in cells in static vs stretched cells and corresponding dSTORM super-resolution images reveal resolved desmosomal plaques in both conditions, with zoomed insets highlighting single desmosomes. (e) Representative linescans and plaque-to-plaque (PtP) distances for static and stretched cells. (f) Quantification of PtP distances reveals a significant increase following stretch (static: 237 ± 35 nm, n=44; stretch: 264 ± 36 nm, n=27; p=0.034). Mann-Whitney U test. Data represents three independent experiments. Normality confirmed by Shapiro-Wilk test.

However, the structural response of DSMs to mechanical forces and the mechanisms through which force is propagated across the DSM remain poorly understood.

Recent advances using super-resolution microscopy, particularly direct stochastic optical reconstruction microscopy (dSTORM), have revealed that DP serves not only as a structural anchor but also as a dynamic regulator of DSM adhesive function. DP exhibits domain-specific conformation during maturation and hyperadhesive transitions, reflecting dynamic adaptations to biochemical cues. dSTORM enables 20 nm resolution imaging of individual DSM plaques not resolvable by conventional microscopy and allows quantitative mapping of plaque-to-plaque distances (PtP). PtP is measured perpendicular to the plasma membrane between the dense cytoplasmic plaques contributed by two adjacent cells. These measurements consistently place DP’s head domain near the membrane, with its tail domain extending into the IDP variably based on rod length and DSM maturation, supporting a model where DP is angled within the plaque (Ainslie et al., 2025; Beggs et al., 2022; Stahley et al., 2016).

Here we examine how mechanical stretch affects DSM architecture in keratinocytes. By integrating biomechanical perturbation with dSTORM, we investigate whether external forces drive the nanoscale organization of DP. We hypothesized that physiological stretch modulates DP tail domain position within the IDP, and this conformational change may enable DSMs to absorb mechanical strain and maintain cell adhesion.

To test this hypothesis, normal human epithelial keratinocytes (NHEKs) were grown on flexible membranes and subjected to 13% uniaxial stretch for 30 minutes or unstretched to serve as control (Fig. 1b-c). The direction (uniaxial) and magnitude of stretch were selected to ensure uniform strain across keratinocytes and to approximate the mid-range of physiological stretch experienced by the epidermis (Gudipaty et al., 2017; Lutz et al., 2020). While held in the stretched position, cells were fixed and immunolabeled with an antibody targeting the DPI rod domain (AA 1900-1950). Widefield fluorescence microscopy revealed diffraction-limited punctate DP staining at cellular borders, corresponding to DSMs, in both groups (Fig. 1d). dSTORM resolved distinct DSM plaque pairs, representing the DPI rod domains contributed by adjacent cells (Fig. 1d-e). To determine whether stretch altered DP architecture, we quantified the distances between the two DPI plaques within individual DSMs. We observed a significant increase in PtP distance in stretched NHEKs compared to static controls (static: 237±35 nm; stretch: 264±36 nm; p=0.034) (Fig. 1f). Because DSM response may vary with orientation relative to the stretch axis, only DSMs with their midline perpendicular to the direction of stretch were included in the analysis.

To further investigate how mechanical stretch influences DSM architecture, we next examined the spatial arrangement of DP’s three splice isoforms: DPI, DPIa, and DPII. These isoforms only differ in length of their central rod domain, and are expressed in the epidermis, with DPI and DPII being the predominant forms. Although their precise functional differences remain unclear, loss of a single isoform has been associated with weakened intracellular adhesion and skin fragility (Cabral et al., 2012).

Because all three isoforms are identical in sequence outside of their rod domain length, current antibodies cannot distinguish between them. Therefore, we used mEGFP-tagged DP constructs stably expressed in DP knockout HaCaT cells, generating separate cell lines for each isoform (Fig. 2a) (Ainslie et al., 2025). This enabled us to independently assess the structural response of each isoform to mechanical stretch. As expected, all DP-mEGFP fusion proteins localized to cell borders (Fig. 2b). Cells were seeded on the flexible membranes and subjected to 13% uniaxial strain for 30 minutes or kept under static conditions. After fixation, cells were labeled with an anti-mEGFP and imaged by dSTORM. In all three cell lines, mechanical stretch resulted in a significant increase in PtP distance, indicating that each isoform undergoes an architectural change following stretch: DPI (static: 191±32 nm; stretch: 210±25 nm; p=0.0322), DPIa (static: 156±22 nm; stretch: 174±18 nm; p=0.0132), and DPII (static: 138±28 nm; stretch: 163±23 nm; p=0.0022). These findings reflect that all DP isoforms undergo stretch-induced remodeling and that mechanosensitivity is an inherent property of DP, regardless of isoform.

**Figure 2.**
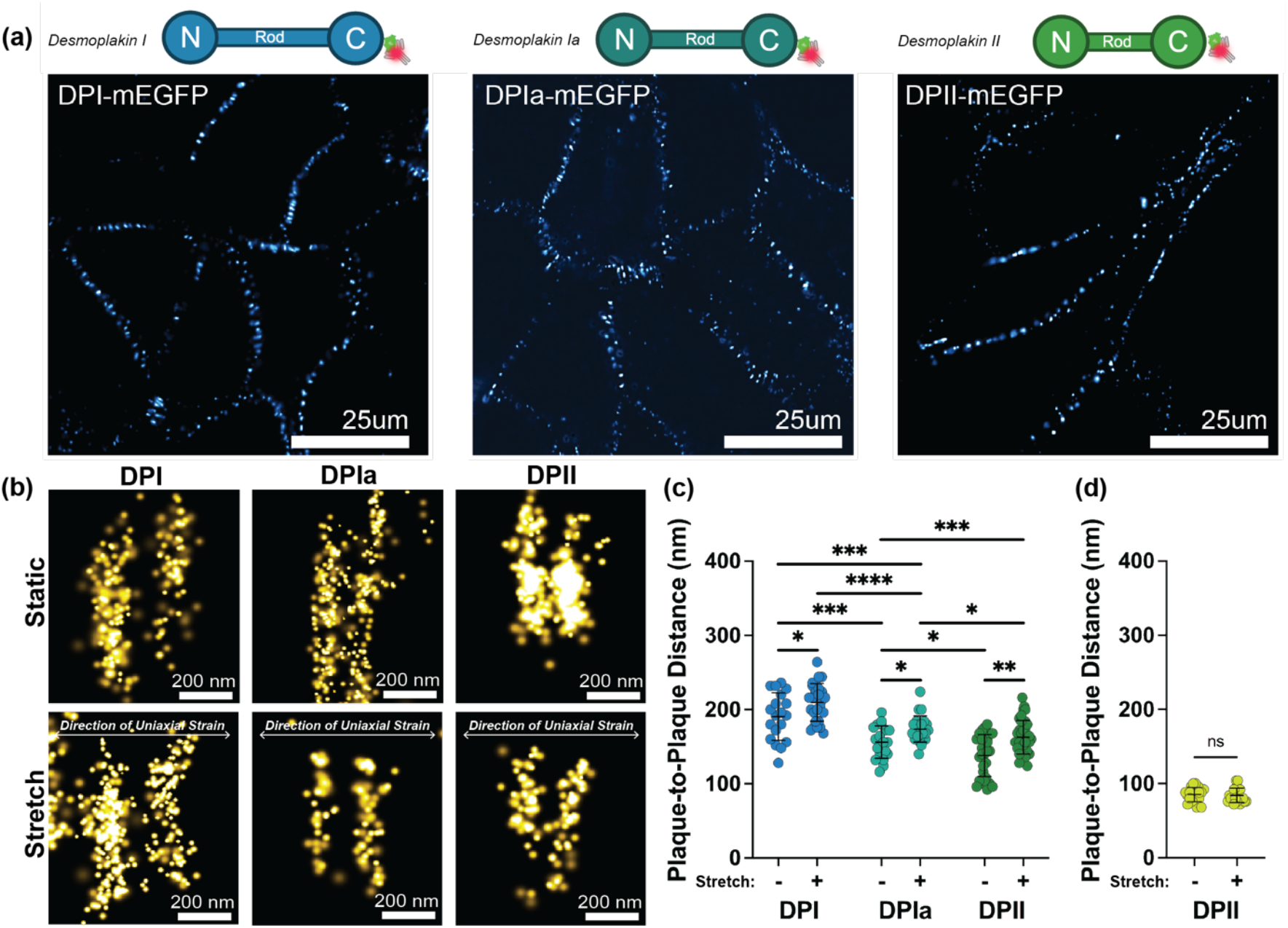
Mechanical stretch induces isoform-specific conformational changes in desmoplakin. (a) Schematics of DP isoforms (DPI, DPIa, and DII) fused to mEGFP above confocal images of DP KO cells stabley expressing each isoform showing DP-mEGFP localization at cell-cell borders. Scale bars = 25 μm. (b) dSTORM super-resolution images of individual desmosomes from cells expressing DPI-, DPIa-, or DPII-mEGFP under static or 13% uniaxial stretch conditions. Scale bars = 200 nm. (c) Quantification of plaque-to-plaque (PtP) distances revealed a significant increase in the spacing between DP C-termini following stretch across all isoforms: DPI (static: 191 ± 32 nm, n = 19; stretch: 210 ± 25 nm, n = 25; p=0.0322), DPIa (static: 156 ± 22 nm, n = 20; stretch: 174 ± 18 nm, n = 23; p=0.0132), and DPII (static: 138 ± 28 nm, n = 29; stretch: 163 ± 23 nm, n = 38; p=0.0022). In contrast, no significant change was observed in PtP distance measured from the DP N-terminus in DPII-mEGFP cells (static: 85 ± 10 nm, n=20; stretch: 84 ± 10 nm, n=20; p=0.5480). Mann-Whitney U test. Data represents three independent experiments. Normality confirmed by Shapiro-Wilk test. Statistical significance is indicated as follows: p < 0.05 (^*^), p < 0.01 (^**^), p < 0.001 (^***^), and p < 0.0001 (^****^); p ≥ 0.05 was considered not significant (ns).

Finally, to test whether the increase in PtP distance reflects a displacement of DP within the plaque, we labeled DPII-mEGFP HaCaT cells with an antibody targeting the DP head domain. PtP measurements showed no significant difference between stretched and static conditions (86±10 nm vs. 80±10 nm, p=0.548, n=20) (Fig. 2d). This suggests that DP does not shift as a unit but rather undergoes conformational or domain-specific architectural changes in response to mechanical stretch.

Compared to static controls, we found that mechanical stretch significantly increased the distance between DP cytoplasmic tails across all isoforms, while the N-terminus domain remained unchanged. This highlights that the tail domain is the primary site of mechanical adaptation with the DSM. These findings provide new insights into the structural dynamics of DP, allowing it to accommodate and dissipate mechanical forces. Importantly, the mechanical responsiveness observed across all isoforms suggests a conserved role for each variant in maintaining DSM adhesion under stress. By adjusting its spatial conformation, rather than shifting as a rigid structure, DP may serve as a molecular buffer, helping to preserve epidermal cohesion and barrier integrity during mechanical challenge. These insights not only advance our understanding of DSM architecture but also have important implications for dermatological health. Defects in DP structure or regulation are implicated in a variety of skin fragility syndromes and blistering disorders. Understanding how DP responds to forces at the nanoscale enhances our ability to interpret how biomechanical dysfunction may compromise the epidermal barrier, informing future strategies for the diagnosis and treatment of skin diseases driven by mechanical stress.

## Ethics Statement

This study did not involve human participants or live vertebrate animals. Primary human keratinocytes were purchased from certified vendors, and desmoplakin isoform cell lines were generated in-house and are available from the corresponding author upon request. All work complied with institutional biosafety and ethical standards.

## Data availability statement

Data is available upon request.

## Acknowledgments

NIH/NIAMS R01 AR072697 and NIH/NIGMS RM1GM145394 to ALM. The UAB High Resolution Imaging Facility was supported by NCI P30 CA013148.

## Author Contributions

Conceptualization: LDS, ALM; Methodology: LDS; Formal Analysis: LDS, CMA; Investigation: LDS, CMA; Writing – Original Draft: LDS; Writing-Review & Editing: MS-K, ALM; Resources: MK-S, ALM; Supervision: ALM; Funding Acquisition: ALM.

## Supplemental Methods

### Cell lines and culture

Four cell lines were investigated: normal human epidermal keratinocytes (NHEKs) (CELLnTEC, Bern, Switzerland) and three mEGFP-tagged isoforms (DPI, DPIa, DPII), each stably expressed DP-knockout in human keratinocytes (HaCaTs). NHEKs were cultured according to the manufacturer’s protocols, and HaCaTs were generated and cultured as described in Ainslie et al., 2025 (Ainslie et al., 2025).

### Extracellular Matrix Coating

Uniflex culture plates (Flexcell International) were coated with Laminin-332 to promote cell adhesion. Laminin-332 (Advanced BioMatrix, Cat. #5463) was diluted to a concentration of 10 μg/mL in 1x DPBS containing Ca^2+^ and Mg^2+^, 3 mL of solution was added to each well, and plates were incubated at 4°C overnight. Prior to cell seeding, the plates were warmed to 37°C, the coating solution was aspirated, and wells were rinsed 3x with DPBS.

### Cell Seeding

Cells were seeded onto the Laminin-332 coated Uniflex plate at a density of 1x10^6^ cells/well. Media was replaced daily and reached 100% confluency after 48 hours and before starting mechanical stretching.

### Flexcell for Cell Stretching

Mechanical strain was applied using the FX-6000™ Tension System (Flexcell, Hillsborough, NC), which utilizes vacuum pressure to deform the flexible-bottomed UniFlex® plates. Arctangle® Loading Stations™ were inserted beneath the culture plates to facilitate uniaxial strain. The system was programmed via FlexSoft® FX-6000™ software to apply a constant uniaxial strain of 13% for a duration of 30 minutes. In order to stain the cells contained in each well of the 6-well dish with more than one antibody, the flexible rubber membrane was cut into 6 square fragments with surgical scissors (Fine Science Tools, 14060-09).

### Fixation and Immunofluorescence

Cells were fixed with 1:1 solution of ice-cold methanol and acetone for 10 mins while FlexCell plates were held in the stretched position at 37°C. Samples were washed three times and blocked with 1% (w/v) Bovine Serum Albumin (BSA), 0.05% Normal Horse Serum (NHS), 0.05% Normal Goat Serum (NGS), and 0.5% Triton X-100. Samples were incubated with primary antibody for 1 hr at 37°C, washed 3x, incubated with secondary antibody for 30 mins at 37°C, washed 3x, and stored at 4°C in PBS.

### Antibodies

NHEKs were imaged using the antibodies anti-desmoplakin (A303-356A Bethyl Laboratories, Montgomery, TX) and anti-rabbit Alexa Fluor 647 (a-21244 Thermo Fisher, Waltham, MA). For detecting mEGFP-tagged DP isoforms in HaCaTs, antibodies with FluoTag-X4 anti-GFP conjugated with Alexa 647 (N0304 NanoTag Biotechnologies, Gottingen, Germany) were used. Primaries were diluted to 1:150, whereas secondaries were used at a 1:1000 dilution.

### Microscopy

Confocal images were obtained on a Ti-2 AXR microscope (Nikon Instruments, Melville, NY) equipped with a 60X 1.42 NA oil immersion objective, and 488 nm laser with Nyquist sampling. dSTORM images were obtained on a Nikon Ti-2 microscope system (Nikon Instruments, Melville, NY) equipped with a 100X 1.49 NA oil immersion objective, 647 nm laser, and Andor iXon EMCCD camera. Each image included 10,000 frames and samples were imaged in a 50mM Tris-HCl, 10mM NaCl, 10% glucose buffer with 5% 1M MEA (Sigma, St. Louis, Missouri) and 2% GLOX (20% 17mg/ml catalase (Roche, Penzberg, Germany) and 14 mg glucose oxidase (Sigma, St. Louis, Missouri) each prepared in 50mM Tris-HCl and 10mM NaCl.

### Image Analysis

Plaque-to-plaque (PtP) distances were quantified using a custom MATLAB-based algorithm as described previously (Beggs et al., 2022). Individual desmosomal plaque pairs were identified in dSTORM reconstructions, and the linear distance between the centroids of adjacent cytoplasmic plaques (representing opposing cell membranes) was measured. Analyses were performed only on DSMs oriented perpendicular to the axis of stretch to ensure consistent spatial interpretation. A minimum of three independent experiments were conducted for each condition. Statistical analyses were performed using Prism 10 (GraphPad Software). Data distribution was assessed by the Shapiro–Wilk test; differences between groups were evaluated by Mann–Whitney U test with significance set at p < 0.05.

